# Solving High-Resolution Forward Problems for Extra- and Intracranial Neurophysiological Recordings Using Boundary Element Fast Multipole Method

**DOI:** 10.1101/567933

**Authors:** Sergey N Makarov, Matti Hämäläinen, Yoshio Okada, Gregory M Noetscher, Jyrki Ahveninen, Aapo Nummenmaa

## Abstract

We present a general numerical approach for solving the forward problem in high-resolution. This approach can be employed in the analysis of noninvasive electroencephalography (EEG) and magnetoencephalography (MEG) as well as invasive electrocorticography (ECoG), stereoencephalography (sEEG), and local field potential (LFP) recordings. The underlying algorithm is our recently developed boundary element fast multipole method (BEM-FMM) that simulates anatomically realistic head models with unprecedented numerical accuracy and speed. This is achieved by utilizing the adjoint double layer formulation and zeroth-order basis functions in conjunction with the FMM acceleration. We present the mathematical formalism in detail and validate the method by applying it to the canonical multilayer sphere problem. The numerical error of BEM-FMM is 2-10 times lower while the computational speed is 1.5–20 times faster than those of the standard first-order FEM. We present four practical case studies: (i) evaluation of the effect of a detailed head model on the accuracy of EEG/MEG forward solution; (ii) demonstration of the ability to accurately calculate the electric potential and the magnetic field in the immediate vicinity of the sources and conductivity boundaries; (iii) computation of the field of a spatially extended cortical equivalent dipole layer; and (iv) taking into account the effect a fontanel for infant EEG source modeling and comparison of the results with a commercially available FEM. In all cases, BEM-FMM provided versatile, fast, and accurate high-resolution modeling of the electromagnetic field and has the potential of becoming a standard tool for modeling both extracranial and intracranial electrophysiological signals.

## 1. Introduction

Electroencephalography (EEG) (Schomer & Lopes da Silva, 2017; Nunes & Srinivasan, 2006) and magnetoencephalography (MEG) (Hämäläinen et al., 1993) record electric potentials and magnetic fields due to neural currents non-invasively. These methods can be used as tools in basic neuroscience, in clinical research, and as diagnostic and monitoring tools in clinical practice. In addition, EEG as well as invasive electrocorticographic (ECoG) recordings can be used in brain-computer interfaces or BCIs (see, e.g., Semprini et al., 2018; Al-Qaysi et al., 2018; Leuthardt et al., 2004) with the goal of mitigating various neurological disabilities (Wolpaw et al., 2002).

Due to the geometry and electrophysiological characteristics of the cortical neurons, a current dipole is used as the elementary source model in the analysis of EEG and MEG (Schomer & Lopes da Silva, 2017; Nunes & Srinivasan, 2006; Hämäläinen et al., 1993; Malmivuo & Plonsey, 1995). Estimation of the distribution of the sources underlying the measured EEG and/or MEG signal patterns, i.e., the solution of the inverse problem, invariably involves comparison of the measured data and those predicted by the sources. Therefore, a sufficiently accurate solution to the electromagnetic forward problem is a prerequisite. Since the EEG/MEG inverse problem is ill-posed, additional constraints are required to render the solution uniqueness (see, e.g., Baillet, Mosher, and Leahy, 2001). In principle, the same procedure applies for the invasive recordings. These recordings contain more detailed spatial information but the locations of the current sources still need to be computationally inferred from the electric/magnetic field data. An additional challenge in the analysis of invasive data is that the set of recording locations is typical sparse and non-uniform.

Several comprehensive and user-friendly open-source software packages for EEG/MEG analysis are currently available, including Brainstorm (Tadel et al., 2011), FieldTrip (Oostenveld et al., 2011), and MNE (Gramfort et al., 2014). However, the corresponding forward-problem solution still offers significant room for improvement. All of the three above-mentioned packages offer the analytical spherically symmetric conductor model and the boundary-element method (BEM) as options for the solution of the forward problem. Additionally, there are “standalone” packages such as Helsinki BEM MATLAB library (Stenroos, Mäntynen, and Nenonen, 2007) and OpenMEEG (Gramfort et al., 2010) that contain the core routines that can be utilized to perform the necessary forward modeling calculations in various cases. The standard boundary element method (BEM) implementations employ three low-resolution layers extracted from the subject’s MRI: scalp, outer skull, and inner skull. The resolution of the scalp, outer skull, and inner skull layers cannot be made very high; it is limited to approximately 7,000 triangles per layer (2,000 default) when Brainstorm is used (Tadel, Bock, and Mosher, 2018). Furthermore, other tissue compartments including cerebrospinal fluid (CSF), gray matter (GM), and white matter (WM) that require a large number of triangular elements for accurate geometric description are not routinely included. As result, significant errors may be generated for the forward problem. This may become increasingly important for the future MEG/EEG modeling, considering the recent technical development of the Optically Pumped Magnetometer (OPM)-based MEG systems (Sander et al., 2012; DARPA, 2018; Iivanainen, Stenroos, and Parkkonen, 2017).

The conventional BEM approach has been extended to include CSF for MEG/EEG forward models (Stenroos & Nummenmaa, 2016). Promising modern techniques include development of a symmetric BEM formulation which significantly improves the accuracy of the BEM method based EEG imaging (Rahmouni, Adrian, Cools, and Andriulli, 2018; Ortiz, Pillain, Rahmouni, and Andriulli, 2018) as well as using a volume integral equation (Rahmouni, Mitharwal, and Andriulli, 2015) to handle anisotropic conductivities in the EEG forward problem (Pillain, Rahmouni, and Andriulli, 2019), which is one major challenge of the boundary element method. However, it is unlikely that the standard BEM approach (Barnard, Duck, and Lynn, 1967; Geselowitz, 1967; Sarvas, 1987; Hämäläinen and Sarvas, 1989; Meijs, Weier, Peters, and van Oosterom, 1989; de Munck, 1992; Hämäläinen et al., 1993; Ferguson, Zhang, and Stroink, 1994; Stenroos, Mäntynen, and Nenonen, 2007; Salinas, Lancaster, and Fox, 2009; Tadel et al., 2011; Stenroos & Sarvas, 2012; Stenroos & Nenonen, 2012; Nummenmaa et al., 2013; Stenroos & Nummenmaa, 2016) could overcome the model size limitation outlined above since its computational burden increases very rapidly with increasing the model resolution. Namely, the BEM matrix is dense with *N*^2^ elements and the direct BEM solution (LU factorization) requires *O* (*N*^3^) operations where *N* is the number of facets. To overcome this issue, the fast multipole method (FMM) has been previously adopted to the MEG/EEG forward problem (Kybic et al., 2005b). Perhaps surprisingly, this approach has not been widely applied for practical MEG/EEG forward modeling problems. The main challenge is to select the optimal BEM formulation and devise an accurate numerical integration scheme for the solution, as well as to couple these with the proper FMM algorithm in an efficient way. Recently, a solution to this problem has been proposed (Makarov, Noetscher, Raij, and Nummenmaa, 2018; Htet et al., 2019) that combines the adjoint double layer formulation of the boundary element method (Barnard, Duck, and Lynn, 1967; Kybic et al., 2005a; Makarov, Noetscher, and Nazarian, 2016; Rahmouni, Adrian, Cools, and Andriulli, 2018) which utilizes surface charges at the boundaries, the zeroth-order (piecewise constant) basis functions, the Galerkin method with accurate near-field integration of the double surface integrals, and the proven FMM accelerator (Gimbutas & Greengard, 2015). This approach does not require the BEM matrix to be formed and inverted explicitly; an iterative solution with *M* iterations requires *O* (*MN*) operations. It has been applied to modeling the transcranial magnetic stimulation (TMS) fields and has demonstrated a fast computational speed and superior accuracy for high-resolution head models as compared to both the standard boundary element method and the finite element method of first order (Makarov, Noetscher, Raij, and Nummenmaa, 2018; Htet et al., 2019). These results have been further confirmed in (Gomez, Dannhauer, Koponen, and Peterchev, 2018).

In this study, the general formalism of the BEM-FMM approach is developed and subsequently applied to numerically challenging forward problem scenarios for electromagnetic recording techniques such as EEG/MEG. The main challenge is to model a large number of internal singular current sources close to tissue conductivity boundaries. The advantage of BEM-FMM is the capability to handle models with accurate conductivity boundaries comprising of up to 50 millions of triangular elements and the generality of the approach including applicability to cases when the conductivity boundaries are not nested and/or may contain “holes” such as skull openings that previously has required a separate formalism to be applied (Stenroos, 2016). The ability of BEM-FMM to overcome these challenges is established with a set of simulations. First, we compare the results given by BEM-FMM with those reported by others authors using the first-order finite element method (Engwer, Vorwerk, Ludewig, and Wolters, 2017; Piastra et al., 2018) in a spherically symmetric case and demonstrate the advantages of our approach. Second, we illustrate the generality and usefulness of our approach in four practical case studies. These examples include: (i) evaluation the effect of a detailed head model on the accuracy of EEG/MEG forward solution; (ii) demonstration of the ability to accurately calculate the electric potential and the magnetic field in the immediate vicinity of the sources and conductivity boundaries; (iii) computation of the field of a spatially extended cortical equivalent dipole layer; and (iv) taking into account the effect a fontanel for infant EEG source modeling and comparison of the results with a commercially available FEM (ANSYS^®^ Maxwell 3D Electromagnetics Suite 2019 R1).

## 2. Materials and Methods

### 2.1 Charge-based formulation of the boundary element method for the secondary field

Two main types of the boundary integral equation formulations for quasistatic modeling are currently being used: the first is framed in terms of electric potential *𝜑*(***r***) while the second is written in terms of electric charge density *ρ*(***r***) at the boundaries (Barnard, Duck, and Lynn, 1967). These two approaches are referred to as the double-layer and the adjoint double-layer formulations, respectively (Rahmouni, Adrian, Cools, and Andriulli, 2018). In addition, there is the “symmetric” formulation (Kybic et al., 2005a; Rahmouni, Adrian, Cools, and Andriulli, 2018), which may have certain computational advantages (Rahmouni, Adrian, Cools, and Andriulli, 2018; Ortiz, Pillain, Rahmouni, and Andriulli, 2018). We chose the adjoint double-layer formulation written in terms of surface charges as a natural foundation for coupling the BEM with the fast multipole method (Rokhlin, 1985; Greengard & Rokhlin, 1987; Gimbutas & Greengard, 2015). Let us consider two (or more) conducting compartments separated by interface *S*. The outer compartment has electric conductivity of *σ*_*out*_ while the inner compartment has conductivity *σ*_*in*_ as shown in Fig. 1a. The vector ***n***(***r***) in Fig. 1 is the outward unit normal vector for the inner compartment. When a primary electric field ***E***^*p*^(***r***, *t*) excitation is applied, surface electric charges with density *ρ*(***r***, *t*) will accumulate at *S* while the electric potential *𝜑*(***r***, *t*) and the normal component of the electric current density remain continuous across the interface. In the quasi-static or low-frequency approximation, the time dependence can be eliminated and therefore is considered as a purely parametric multiplicative factor.

**Fig. 1.**
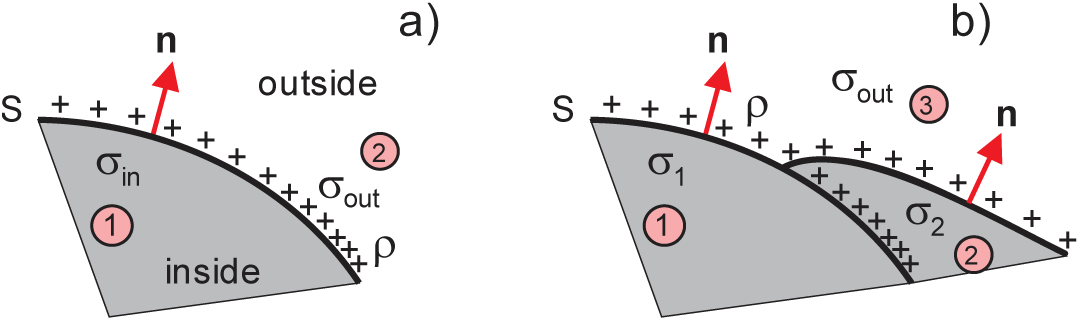
Boundary between two conducting compartments with different conductivities and surface charge density *ρ*(***r***) residing at the boundary.

The most widely used potential-based approach results in the integral equation (Barnard, Duck, and Lynn, 1967; Geselowitz, 1967; Sarvas, 1987; Hämäläinen and Sarvas, 1989; Meijs, Weier, Peters, and van Oosterom, 1989; de Munck 1992; Hämäläinen et al., 1993)

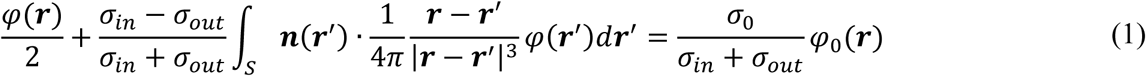

where *σ*_0_ = 1 S/m is the unit conductivity and *𝜑*_0_(***r***) is the potential arising from a given excitation in a homogeneous medium with unit conductivity. For EEG/MEG applications, it is the electric potential of a current dipole. Eq. (1) is well suited for EEG studies since it gives us the solution directly in the form of the electric potential on the scalp surface as well as on the other interfaces between tissue compartments with different conductivities. It is also well suited for MEG studies since, after solving the surface potentials, the magnetic field can be found using Geselowitz’ formula (see, e.g., Sarvas, 1987). However, Eq. (1) is derived using Green’s second identity (Hämäläinen et al., 1993) and thus only valid for closed surfaces with one value of external conductivity, in particular for surfaces enclosed into each other in the form of a nested structure. Inclusion of surface junctions (*e.g.*, an opening in a skull sketched in Fig. 1b) requires a special treatment (Stenroos, 2016).

On the other hand, the surface-charge approach uses the boundary conditions only and therefore directly applies to a wider range of volume conductor geometries. The corresponding integral equation is obtained by writing the total electric field ***E***^*t*^ in a form that considers the primary field ***E***^*p*^ and a conservative contribution of the secondary induced surface charge density ***E***^*s*^:

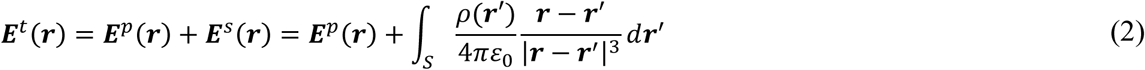

where *ɛ*_0_ is the permittivity of vacuum. Taking the limit of Eq. (2) as ***r*** approaches surface *S* from both sides and using the continuity condition for the normal current component, *σ****E***^*t*^(***r***), one obtains the adjoint double-layer equation ((Barnard, Duck, and Lynn, 1967; Makarov, Noetscher, and Nazarian, 2016; Rahmouni, Adrian, Cools, and Andriulli, 2018):

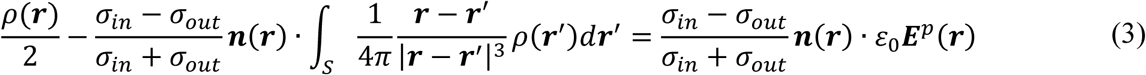

Eq. (3) is directly applicable to the surface junction case from Fig. 1b. In contrast to Eq. (1), the normal-vector multiplication becomes external in the adjoint operator, i.e., it appears outside of the potential integral. This leads to computing the gradient of a single layer in Eq. (3) instead of the potential of the double layer in Eq. (1).

### 2.2. Fast multipole method (FMM) for the secondary field

The fast multipole method (Rokhlin, 1985; Greengard & Rokhlin, 1987) speeds up computation of a matrix-vector product by many orders of magnitude. Such a matrix-vector product appears when an electric field from many point sources *ρ*(***r***^′)^ in space has to be computed at many observation points ***r***. In the present problem, this computational task emerges from the discretization of the surface integral in Eq. (2) or in Eq. (3). Assuming zeroth-order piecewise constant basis functions (pulse bases), one has

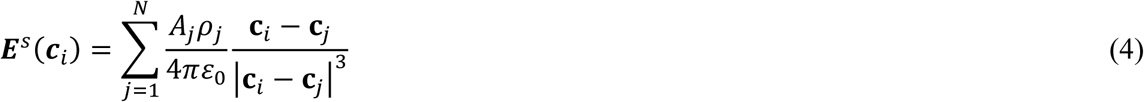

where *A*_*i*_, ***c***_*i*_, *i* = 1, …, *N* are the areas and centers of the triangular surface facets *t*_*i*_ and *ρ*_*i*_ are the triangle surface charge densities. An approximation of the expression on the right-hand side of (4) is computed via the FMM with drastic improvements in the computational speed. We adopt, integrate, and use an efficient and proven version of the FMM (Gimbutas & Greengard, 2015) originating from its inventors. In this version, there is no *a priori* limit on the number of levels of the FMM tree, although after about thirty levels, there may be floating point issues (L. Greengard, private communication). The required number of levels is determined by a maximum permissible least-squares error or method tolerance, which is specified by the user. The FMM is a FORTAN 90/95 program compiled for MATLAB. The tolerance level `iprec` of the FMM algorithm is set at 0 or 1 (the relative least-squares error is guaranteed not to exceed 0.5% or 0.05%, respectively). This FMM version allows for a straightforward inclusion of a controlled number of analytical neighbor integrals to be precisely evaluated as specified below.

### 2.3. Correction of neighboring terms and iterative solution for secondary field

Approximation (4) is inaccurate for the neighbor facets. Using Galerkin (or strictly speaking Petrov-Galerkin) method with the same pulse bases as testing functions, we employ a refined computation:

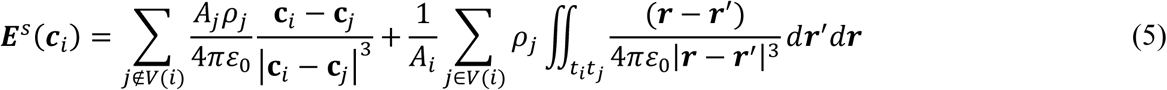

where *V*(*i*) is a neighborhood of observation triangle *t*_*i*_. The inner integrals in Eq. (5) are computed analytically (Wilton et al., 1984; Wang et al., 2003; Makarov, Noetscher, and Nazarian, 2016) while the outer integrals use a Gaussian quadrature on triangles of 10^th^ degree of accuracy (Cools, 2003). We have implemented two methods for the selection of *V*(*i*): a ball neighborhood with a radius *R* such that |***c***_*i*_ − ***c***_*j*_| ≤ *R* and a neighborhood of *K* ≪ *N* nearest triangular facets; they yield similar results. However, the second method leads to fixed-size arrays of pre-calculated double surface integrals and to a faster speed, making it the preferred method. Inclusion of a small number of precomputed neighbor integrals (three to sixteen) drastically improves the convergence of the iterative solution; inclusion of a larger number has a negligible effect. The present BEM-FMM approach performs precise analytical integration over *K* = 16 closest neighbor facets. Eq. (3) is then solved iteratively using GMRES (generalized minimum residual method (Barrett et al., 1994; Saad, 2003) implemented by Drs. P. Quillen and Z. Hoffnung of MathWorks, Inc. Its overall performance and convergence are excellent, especially for complicated head geometries.

After the solution for the surface charges is obtained, the condition of an electrically neutral system is explicitly enforced, with the total charge equal to zero, i.e.,

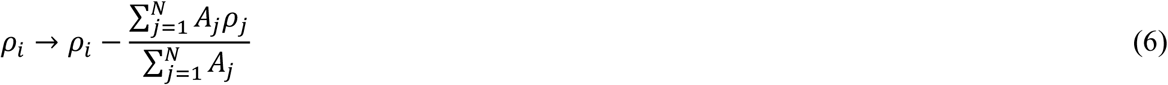

This correction has a negligible effect on the resulting electric/magnetic fields but improves solution accuracy for the electric potential, which is defined up to an additive constant. When the charge conservation law is introduced, this constant no longer needs to be adjusted.

### 2.4. Fast multipole method (FMM) for the primary field

A finite-length EEG/MEG dipole (Nunes & Srinivasan, 2006; Hämäläinen et al, 1993; Malmivuo & Plonsey, 1995) shown in Fig. 2 is constructed from a (isotropic) current source of strength *I*_0_ and electric potential *𝜑*_+_(***r***) = *I*_0_/4*πσ*|***r*** − ***p***_1_| at ***p***_1_ and a current sink −*I*_0_ with electric potential *𝜑*_−_(***r***) = − *I*_0_⁄4**π*σ*|***r*** − ***p***_2_| at ***p***_2_ as shown in Fig. 2. The surrounding medium has conductivity *σ*. The primary electric potential is thus

**Fig. 2.**
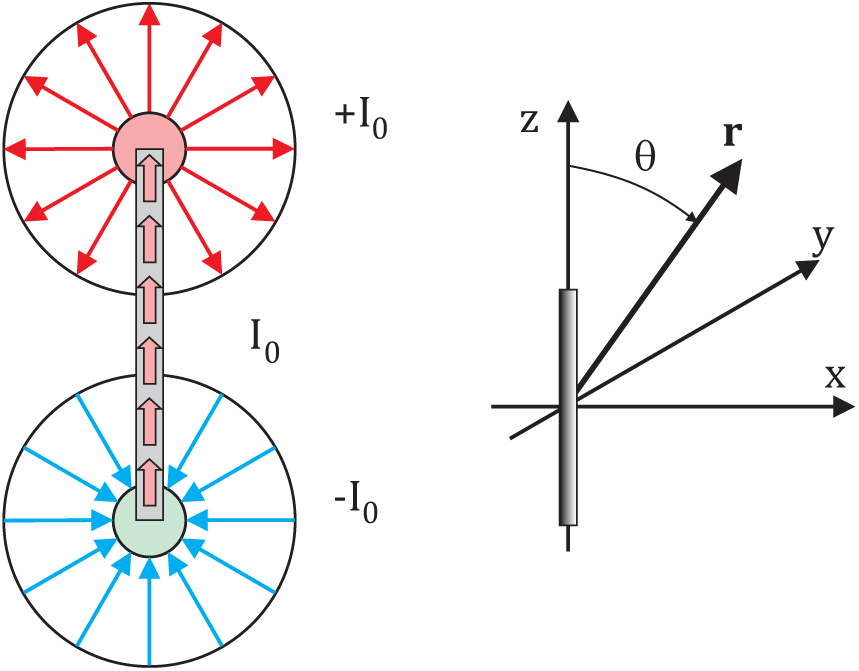
Finite length EEG/MEG dipole.

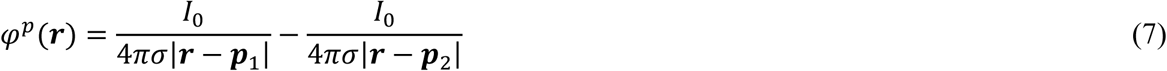

When the vector distance ***d*** = ***p***_1_ − ***p***_2_ from the sink to the source approaches zero, one has the well-known point-dipole expression *𝜑*^*p*^ = ***Q*** ∙ ∇(1⁄|***r*** − ***r***_1_|)⁄4*πσ* with ***Q*** = *I*_0_***d*** being the vector dipole moment (in A⋅m). The primary electric field is

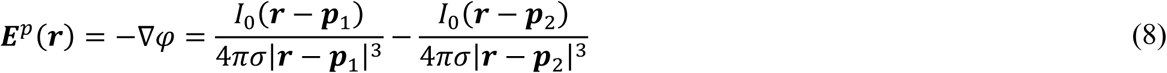

The magnetic vector potential, ***A***, of any point current source (or their linear combination) is a conservative field (Balanis, 2012). Therefore, its curl is zero so that the magnetic flux, ***B*** = ∇ × ***A***, for the combination of two isotropic current sources (7) vanishes everywhere in space.

Both *𝜑*^*p*^ and ***E***^*p*^ generated by the sources located at ***p***_*j*_ are computed at a large number of target points ***r***_*i*_. The target points may be chosen anywhere in the volume or they may coincide with the triangle centers ***c***_*i*_. In either case, we directly apply function lfmm3dpart from the FMMLIB3 library of the fast multipole method (Gimbutas & Greengard, 2015) to all current sources and to all required target points. This gives us the potential *𝜑*^*p*^(***r***_*i*_) and the field ***E***^*p*^(***r***_*i*_).

When the targets are ***c***_*i*_, a ball neighborhood *V*(*i*) with a radius *R* such that |***c***_*i*_ − ***p***_*j*_| ≤ *R* is introduced. The typical *R* is 5-10 times the average edge length. If the source ***p***_*j*_ is within this sphere, its potential and field contributions are corrected by replacing the center-point approximation by accurate integration (the current source of strength *I*_0_ is assumed below) over the triangle area,

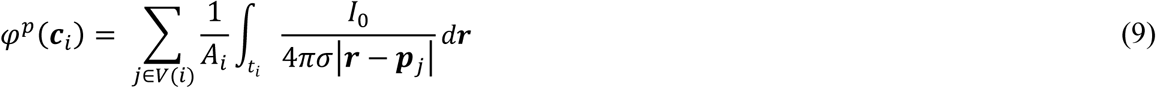

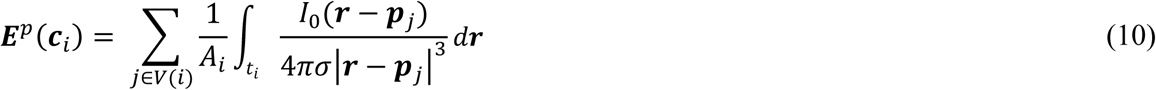

In other words, we compute the primary field by accurate averaging over the entire triangle area if the source is close to the triangle, which may become a decisive advantage for certain situations. The integrals over triangular facets in Eqs. (9), (10) are computed analytically (Wilton et al., 1984; Wang et al., 2003; Makarov, Noetscher, and Nazarian, 2016).

For MEG dipoles, we additionally consider a filamentary current between the two current sources in Fig. 2. Such a current does not give a contribution to the conservative electric field (7),(8), but it creates a solenoidal magnetic vector potential ***A*** and the respective primary magnetic field ***B***^*p*^ = ∇ × ***A***. For a small straight element of current *I*_0_ with length *d* and center ***p***_*j*_, which has a unit direction vector ***n***_*j*_

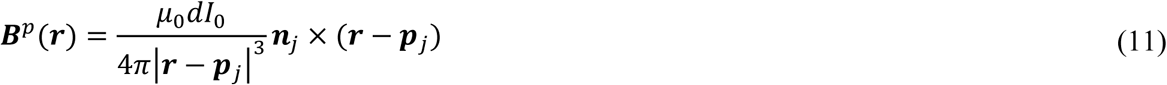

where ***r*** is the target point and *μ*_0_ is the permeability of vacuum. Eq. (11) cannot be evaluated using the FMM directly for multiple source and target points. However, since ***n***_*j*_ × (***r*** − *p*_*j*_) ⋅ ***e***_*k*_ = (***e***_*k*_ × ***n***_*j*_) ⋅ (***r*** − ***p***_*j*_), where ***e***_*k*_ = ***e***_{*x*,*y*,*z*}_ are the three orthogonal unit vectors, we find

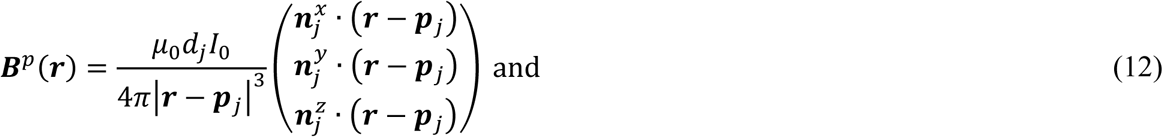

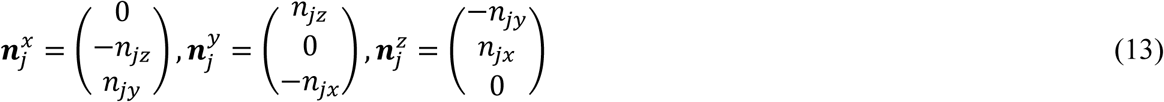

which is equivalent to the electric potential of a double layer (layer of electric dipoles) to be computed three times and with the three different sets of the “direction vectors” given in Eqs. (13). This task is thus accomplished by applying function `lfmm3dpart` from the FMMLIB3 library of the fast multipole method (Gimbutas & Greengard, 2015) three times. Since the magnetic field is measured outside the cortical volume, no refinement of the FMM result is necessary.

### 2.5. Fast multipole method for the total potential (EEG) and total magnetic field (MEG)

After the electric charge density *ρ*_*j*_ for every *j*-th triangular facet at the conductivity boundaries is computed via the iterative solution, the secondary electric potential *𝜑*^*s*^ for every *i*-th triangular facet is found from the familiar electrostatic expression using the FMM. We apply function `lfmm3dpart` from the FMMLIB3 library of the fast multipole method (Gimbutas & Greengard, 2015) and ignore the self-term. This formulation is again inaccurate when the *i*-th and *j*-th triangles are close to each other or coincide. Therefore, it is corrected similar to Eqs. (9), (10), that is

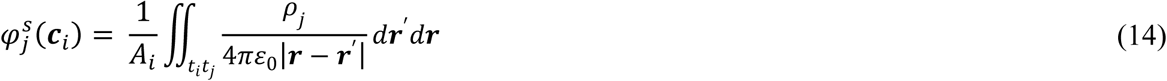

Inner integrals in Eq. (14) are computed analytically (Wilton et al., 1984; Wang et al., 2003; Makarov, Noetscher, and Nazarian, 2016); the outer integrals use a Gaussian quadrature of 10^th^ degree of accuracy (Cools, 2003). For the neighborhood of *K* = 16 closest facets, the double surface integrals (14) are precomputed and then stored in memory to enable fast repetitive postprocessing.

After the secondary electric potential *𝜑*^*s*^ is obtained, the continuous secondary magnetic field ***B***^*s*^ of volumetric currents caused by surface charges in the conducting medium is found using Stokes theorem (Hämäläinen et al, 1993), which is Geselowitz’ formula

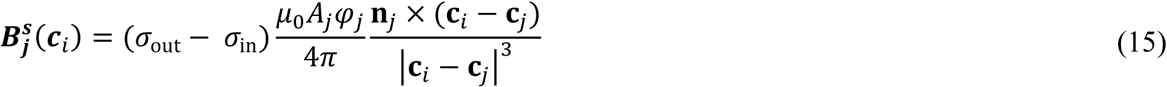

To compute the sum of contributions (15) for all source triangles and for all required target points, we apply the FMM approach three times, following the transform introduced in Eqs. (12) and (13), respectively. We also apply corrections conceptually identical to Eqs. (5),(9),(10),(14) for target points close to the conductivity boundaries or directly on the conductivity boundaries.

## 3. Results

### 3.1. Accuracy of BEM-FMM in the sphere model

We test the accuracy and speed of the proposed BEM-FMM numerical solver against previously published first-order FEM solutions (continuous Galerkin or CG and discontinuous Galerkin or DG) for EEG and MEG problems (Engwer, Vorwerk, Ludewig, and Wolters, 2017; Piastra et al., 2018). We replicate the benchmark tests using our approach and use the published results as a reference. To this end, a four-layer sphere model is created for which well-known analytical solutions for EEG (Zhang 1995; Mosher, Leahy, and Lewis, 1999) and MEG (Sarvas, 1987) exist. Fig. 3a shows the problem geometry and conductivity values previously employed (Engwer, Vorwerk, Ludewig, and Wolters, 2017; Piastra et al., 2018).

**Fig. 3.**
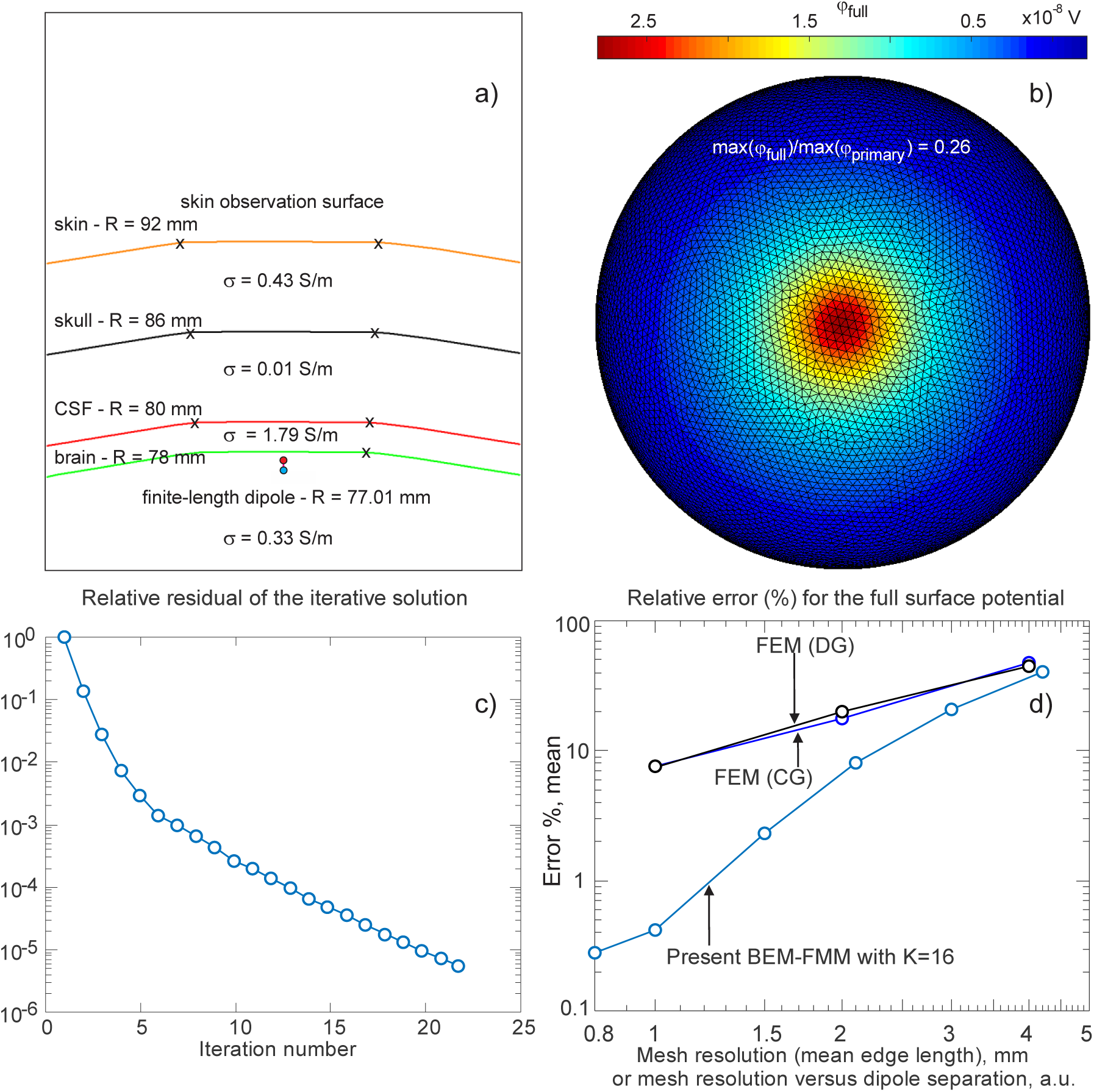
a) Geometry of the sphere model; b) The potential distribution at the skin surface for the combined mesh with the resolution (mean edge length) approximately three times the dipole separation distance. c) Convergence speed of the BEM-FMM solution; d) Relative errors of the FEM and BEM-FMM methods as functions of the mesh resolution.

Since computation of the fields of dipoles located close to the CSF boundary poses highest numerical challenges, we used a source located 0.99 mm inside the CSF boundary (eccentricity 0.987 according to Engwer, Vorwerk, Ludewig, and Wolters, 2017; Piastra et al., 2018). We created six surface sphere meshes with the number of facets ranging from 6,000 to 400,000 using a high-quality surface mesh generator (Persson and Strang, 2004; Persson, 2005). The four boundaries were obtained from a prototype mesh by scaling. The total mesh size thus ranged from 24,000 to 1,600,000.

The EEG problem is considered first. A 0.1 mm long vertical electric dipole schematically shown in Fig. 3a and separated by 0.99 mm from the CSF boundary is employed. The dipole length was chosen to be much smaller than the distance to the closest conductivity boundary (CSF) to be consistent with the analytical solution that uses the point-dipole model (Zhang, 1995; Mosher, Leahy, and Lewis, 1999). The dipole moment *Q* = 1 nA ∙ m. Fig. 3b shows the corresponding potential distribution at the outermost sphere (“skin”) surface for the combined mesh with the segmentation resolution (mean edge length) approximately three times the dipole separation distance and with about 0.05 M facets in total. Fig. 3c shows the convergence speed of the BEM-FMM solution. The final relative residual is required to be less than 5 ⋅ 10^−6^. Next, 300 random radial dipole positions at the same eccentricity of 0.987 within the “brain” sphere surface are generated. An average relative error between numerical and analytical solutions is computed for all tested sphere meshes exactly following (Engwer, Vorwerk, Ludewig, and Wolters, 2017), i.e.

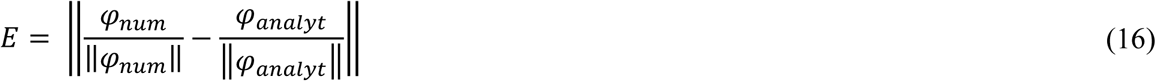

where ‖∙‖ is the 2-norm of the vector of potential values at the skin surface. Fig. 3d shows *E* as a function of surface mesh resolution together with the corresponding result from (Engwer, Vorwerk, Ludewig, and Wolters, 2017), see Figure 7 in the original. These data are for the continuous and discontinuous finite-element Galerkin method, respectively, and for the identical mesh resolutions but achieved with the hexahedral volumetric FEM meshes. Fig. 3d indicates that the BEM-FMM generates a smaller error for all considered mesh resolutions and that its accuracy may exceed the FEM accuracy reported in Ref. (Engwer, Vorwerk, Ludewig, and Wolters, 2017) by an order of magnitude at resolutions on the order of the dipole separation distance.

The MEG problem from Ref. (Piastra et al., 2018) is considered next. The volume conductor geometry of the benchmark test remains unchanged but the MEG test source is a 0.1 mm long tangential current dipole schematically shown in Fig. 4a and separated by 0.99 mm from the CSF boundary (with the same eccentricity factor of 0.987). Following (Piastra et al., 2018), the magnetic field is evaluated on a sphere surface with the radius of 110 mm also shown in Fig. 4a. Fig. 4b shows the magnitude distribution of the magnetic field at this surface for the combined mesh with segmentation resolution approximately three times the dipole separation distance. Fig. 4c shows the convergence speed of the corresponding BEM-FMM solution as a function of the iteration number. The final relative residual is required to be less than 5 × 10^−6^.

**Fig. 4.**
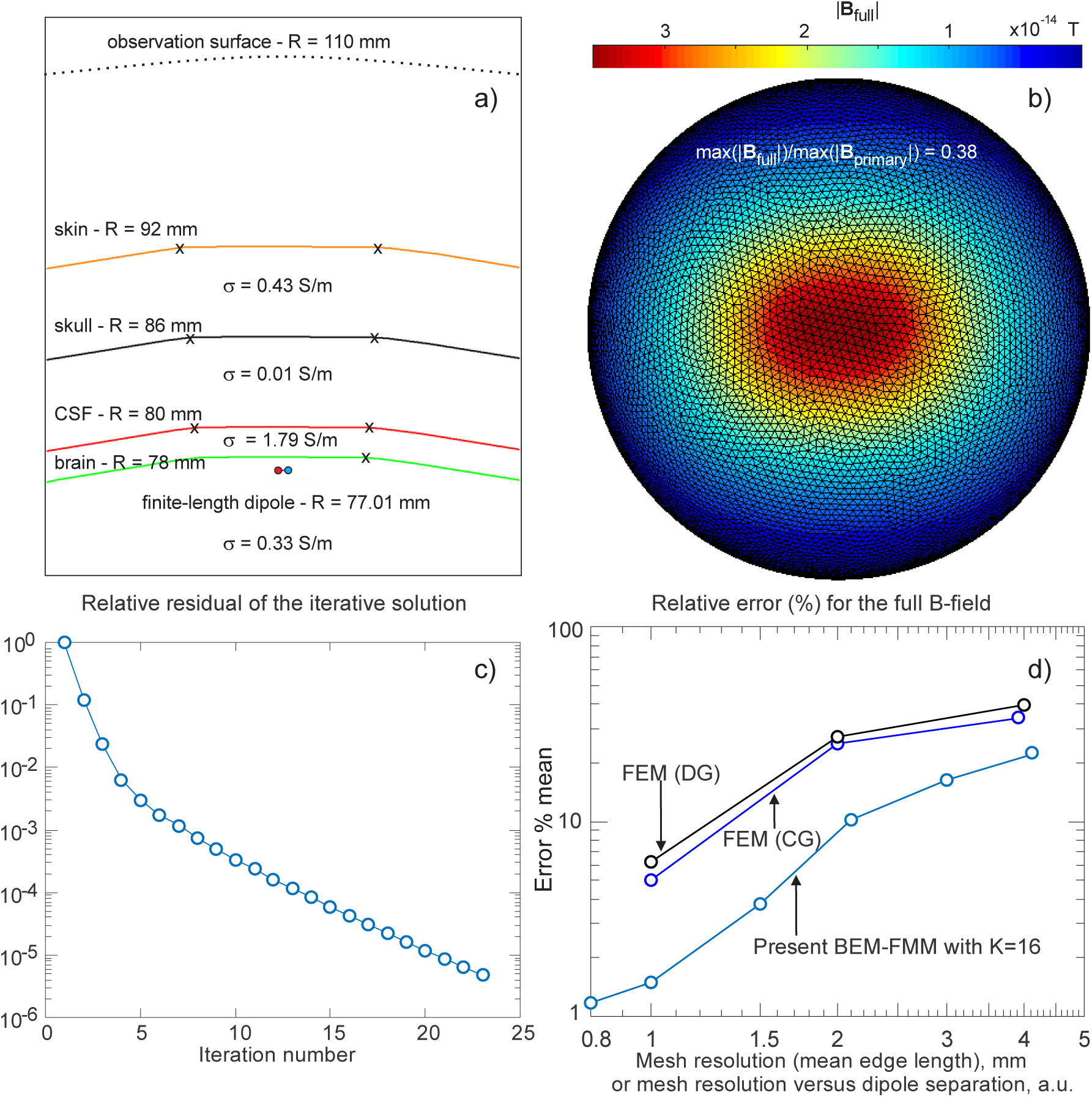
a) Geometry of a multilayered isotropic sphere; b) Distribution of the amplitude of the magnetic field at the observation surface for the combined mesh with the resolution (mean edge length) approximately three times the dipole separation distance; c) Convergence speed of the BEM-FMM solution; d) Relative errors of the FEM and BEM-FMM methods as functions of the mesh resolution.

Next, 300 random tangential dipole positions/directions at the same eccentricity of 0.987 within the “brain” sphere surface are generated. An average relative error between numerical and analytical solutions is computed for all tested sphere meshes exactly following (Piastra et al., 2018), i.e.

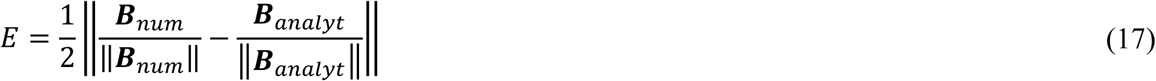

where 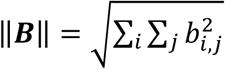 is the Frobenius or *L*_2,2_ norm of the matrix of magnetic field values at the observation surface. Fig. 3d shows *E* as a function of surface mesh resolution together with the corresponding result from Ref. (Piastra et al., 2018), see Figure 8 in the original, for continuous and discontinuous FGEM Galerkin method, and for identical mesh resolution but achieved with hexahedral volumetric meshes. Fig. 4d indicates that the BEM-FMM generates a smaller error for all considered mesh resolutions and that its accuracy exceeds the FEM field accuracy reported in (Piastra et al., 2018) by a factor of 1.5–4.

### 3.2. Computation speed in the sphere model

For the EEG problem shown in Fig. 3, Ref. (Engwer, Vorwerk, Ludewig, and Wolters, 2017) reports a comprehensive evaluation of computational time for the first-order finite-element methods with non-uniform meshing, which will be used for comparison. Either a single dipole source or a group of 4724 dipoles were been considered. Based on these data, Table 1 compares the method speed given the nearly identical mesh resolution close to the conductivity boundaries. The present BEM-FMM solution again uses the relative residual of 5 ⋅ 10^−6^. Table 1 reports computational performance for 1 and 2 mm mesh resolutions. There was no effort to parallelize either of the methods (DUNE FEM software described in online Ref. (DUNE software) used by Engwer, Vorwerk, Ludewig, and Wolters, 2017) or present BEM-FMM running within MATLAB platform with major routines compiled in FORTRAN). However, both software packages (DUNE and core MATLAB) employ multithreading for linear algebra operations, allowing them to execute faster on multicore-enabled machines.

**Table 1.** EEG computation run times for a single source and for 4724 dipole sources including the right-hand side (RHS) computations and compared to Ref. (Engwer, Vorwerk, Ludewig, and Wolters, 2017) labeled as (*). Mesh sizes of 2 mm mesh (0.19 M triangles/0.5 M hexahedra) and 1 mm mesh (0.83 M triangles/3.9 M hexahedra) were used. The 4724 vertical dipoles were randomly located at the distance of 0.99 mm from the CSF boundary.

| Method | Mesh resolution [mm] | Computer Hardware/Software | Run time for a single source [sec] | Run time for 4724 sources including RHS comput. [sec] |
| --- | --- | --- | --- | --- |
| CG FEM (*) | 2 | Intel Xeon E5-2698 v3 CPU (2.30 GHz) Linux | 20 | 42810 |
| DG FEM (*) | 2 | Intel Xeon E5-2698 v3 CPU (2.30 GHz) Linux | 136 | 52689 |
| BEM-FMM | 2 | Intel Xeon E5-2698 v4 CPU (2.20 GHz) MATLAB 2018a, Windows | 19 (0.9 per iteration) | 34 |
| CG FEM (*) | 1 | Intel Xeon E5-2698 v3 CPU (2.30 GHz) Linux | 185 | 336339 |
| DG FEM (*) | 1 | Intel Xeon E5-2698 v3 CPU (2.30 GHz) Linux | 1468 | 452112 |
| BEM-FMM | 1 | Intel Xeon E5-2698 v4 CPU (2.20 GHz) MATLAB 2018a, Windows | 77 (3.5 per iteration) | 92 |

Table 1 shows that, when the right-hand side computational time is excluded, the speed of the BEM-FMM algorithm for a single-dipole is about the same as the speed of the standard FEM (CG) for the coarse 2 mm mesh and exceeds the FEM speed for the fine 1 mm mesh. We emphasize that our platform is based on MATLAB^®^ and that we use complex arithmetic of the FMMLIB3 library of the fast multipole method (Gimbutas & Greengard, 2015). Therefore, we would expect a further speed improvement by approximately a factor of 2 when using a real-valued version of the FMM currently under development.

The results are even more favorable for multiple EEG dipoles and for the corresponding right-hand side computations. One main challenge in applying the finite-element methods to solve the EEG/MEG forward problem is to deal with the strong singularity at the current dipole source. Different singularity extraction approaches have been proposed; among them the subtraction approach developed in (Wolters et al., 2007; Engwer, Vorwerk, Ludewig, and Wolters, 2017; Piastra et al., 2018) and reported in Table 1. This approach very significantly increases the overall FEM run time as Table 1 shows. On the other hand, the BEM-FMM does not need to perform the singularity extraction since there are no volumetric elements containing the sources themselves. Strong and rapidly varying near fields for a small number of nearby surface facets are handled by analytical surface integration in Eqs. (5),(9),(10),(14). As a result, the fast multipole method requires about a minute of run time for the problem of the same complexity and using nearly the same computer hardware. Since the native FMM is very fast for source field calculations, the number of sources in the present approach is practically unlimited, which is of importance for EEG/MEG distributed inverse problem approaches where the entire discretized cortex is employed as the space of possible sources. This is illustrated in Table 1 where we report a very modest (approximately 15 sec) increase in the computation time for 4724 sources.

### 3.3. Effect of anatomically detailed modeling

A triangular surface-based head model for subject #101309 from the Population Head Model Repository (Lee et al., 2016; IT’IS Foundation, 2017) originated from Connectome Project (Van Essen et al., 2012) has been solved via BEM-FMM. The model includes the following seven compartments: cerebellum, CSF, GM, skin (or scalp), skull, ventricles, and WM, and results in 700,000 triangular facets in total. The average mesh resolution (edge length) is 1.5 mm for scalp and skull and 1.1 mm for CSF, GM, and WM. Tissue conductivities are adopted from the IT’IS database (Hasgall et al., 2018) with the average scalp conductivity chosen as 0.333 S/m. With the BEM-FMM approach, an EEG forward computation executes in less than 2 minutes on an ordinary server.

To demonstrate the importance of the anatomical detail and accuracy of the head model, a single finite-length (1.8 mm) cortical dipole with the moment *Q* = 1.8 nA ∙ m is placed in the vicinity of the right motor cortex at two different orientations as shown in Fig. 5a,b.

**Fig. 5.**
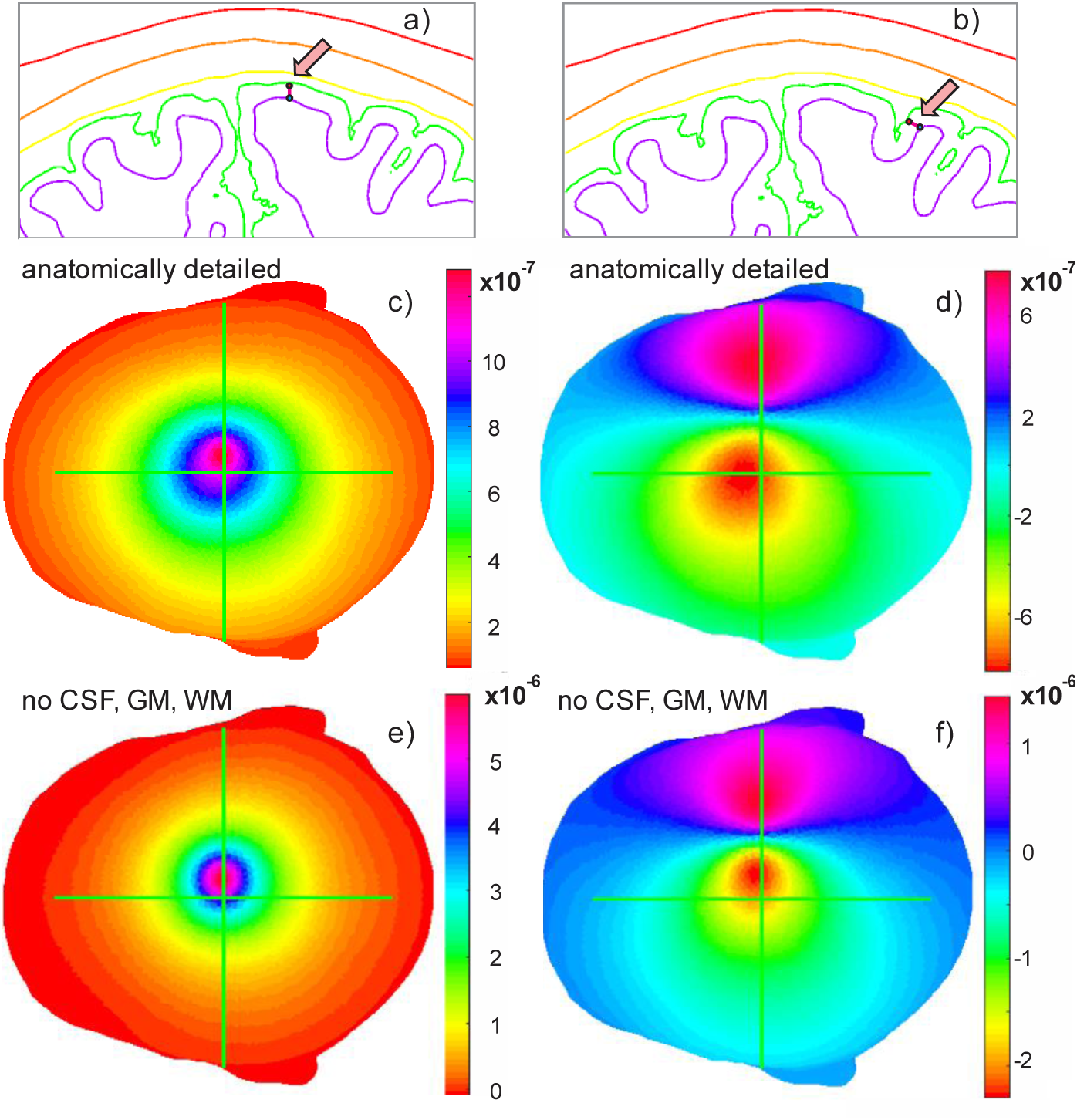
Electric potential distribution in V over the scalp surface for nearly radial (a) and nearly tangential (b) cortical dipoles (*Q* = 1.8 nA ∙ m) at the primary motor cortex (coronal plane); c), d) Solutions with all brain compartments included; e), f) Approximate solutions with only three brain compartments (scalp, inner skull, outer skull) included. Note the differences in the maximum amplitudes.

The scalp potential was computed with all brain compartments present and with only three brain compartments present, respectively, which is similar to OpenMEEG software (Gramfort et al., 2010; Tadel, Bock, and Mosher, 2018). There are clear differences in the scalp potential distribution (Fig. 5 c-f), especially in the amplitude. Quantitative measures are given in Table 2. They are the relative difference measure (RDM) or topographical error given by RDM = ‖*𝜑*_1_/‖*𝜑*_1_‖ − *𝜑*_2_/‖*𝜑*_2_‖‖ and the logarithmic magnitude (lnMAG) error given by lnMAG = ln(‖*𝜑*_1_‖/‖*𝜑*_2_‖). The topographical error in Table 2 may be as large as 67%.

**Table 2.**
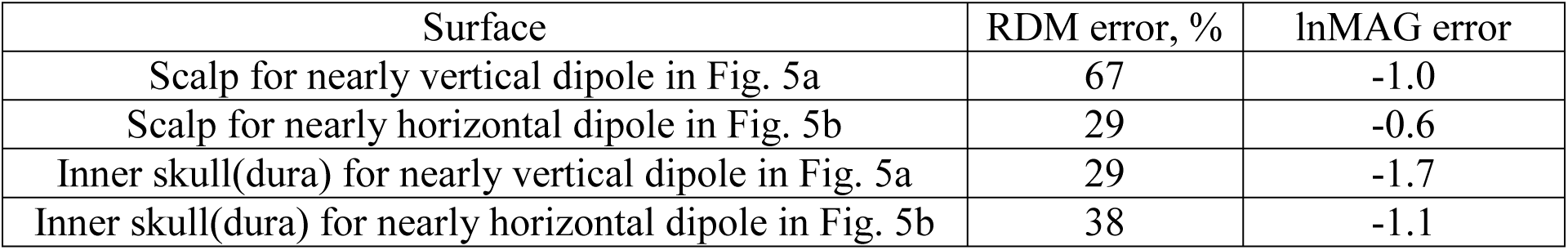
Topographical error and magnitude error for the electric potential between the complete model and that omitting the CSF, WM, and WM interfaces.

### 3.4. High-resolution modeling of intracranial fields

In addition to the extracranial EEG/MEG fields, the BEM-FMM method can be utilized to model the intracranial fields in the immediate vicinity of the neuronal sources. This can be beneficial for analyzing ECoG and/or LFP data and can help us to understand the relationships between intra-and extracranial fields. The surface mesh from the previous example was refined using a 1×4 barycentric triangle subdivision and then surface-preserving Laplacian smoothing (Vollmer, Mencl, and Müller, 1999a,b) was applied. This results in a model that has an average mesh resolution of 0.7 mm (0.6 mm for GM and WM) and 2.8 M triangular facets in total. This high-resolution model runs reasonably fast, in approximately 200 seconds in Windows MATLAB environment given 14 iterations and the relative residual below 10^−4^.

A 1.8 mm long and nearly radial dipole with the moment *Q* = 1 nA · m, directed outwards is placed near the primary motor cortex as shown in Fig. 6a in both coronal and sagittal planes.

Figs. 6c, d show the resulting volume fields (electric and magnetic) in the two planes. Every field plot is based on 250,000 observation points at a spacing of 0.1 mm. The total electric field magnitude is given in the coronal plane in Fig. 6c while Fig 6d shows the absolute value of the dominant tangential component *B*_*x*_ in the sagittal plane. In order to display large field variations, we use the decibel scale as explained in the respective figures. The electric field is indeed observed to be discontinuous at the interfaces between the tissue compartments while the magnetic field is not. Fig. 6c clearly demonstrates how the electric field of the current dipole is attenuated and smeared by the scalp; both the amplitude and the spatial specificity of the recordings would greatly increase if the electrodes could be attached to the skull instead of skin. On the other hand, Fig. 6d illustrates a magnetic-field lacuna that is typical for a nearly radial dipole (which in the spherically symmetric model would produce no extracranial field). However, this lacuna is bent and is shifted quite significantly due to the intricate geometry of the anatomically realistic conductivity boundaries resulting in diminished yet non-zero field outside the head.

**Fig. 6.**
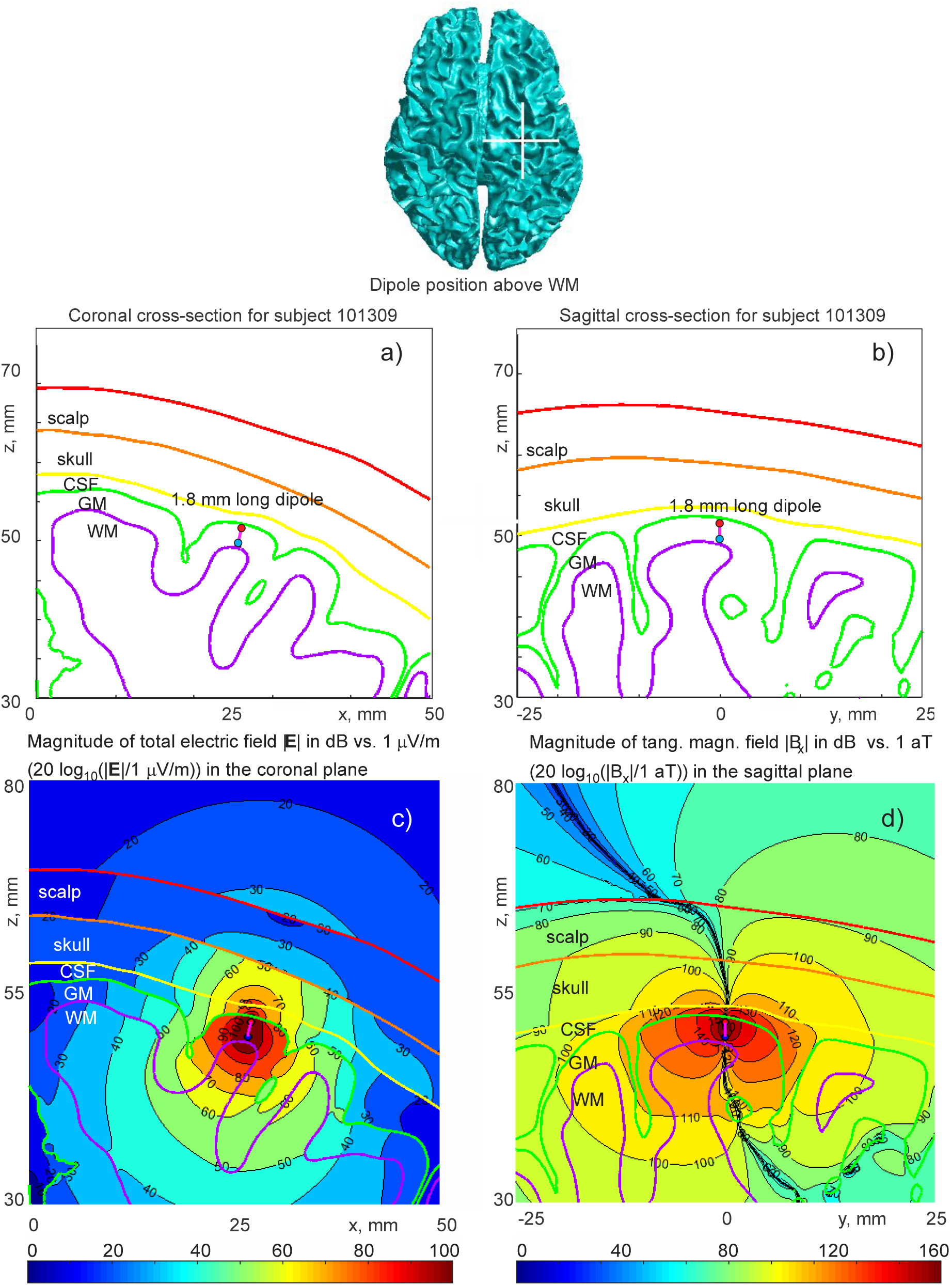
Single-dipole EEG/MEG distributions computed with the BEM-FMM approach. a), b) Problem geometry; c) The electric field magnitude in the coronal plane; d) The absolute value of dominant tangential component *B*_*x*_ in the sagittal plane.

Fig. 7 is the zoomed in version of Fig. 6c; it shows the electric field magnitude distribution in the immediate vicinity of the EEG dipole in Fig. 6c. The same logarithmic scale is used. For every observation point in Fig. 6 or in Fig. 7, the ball neighborhood with the radius *R* = 5*l*, where *l* is average edge length, is introduced. If a surface triangle in question belongs to this domain, the corresponding charge contribution into the total electric field is computed by analytically evaluating the field integral in Eq. (2). Otherwise, the central-point approximation is used which implies that the distributed charge is concentrated at the triangle center.

**Fig. 7.**
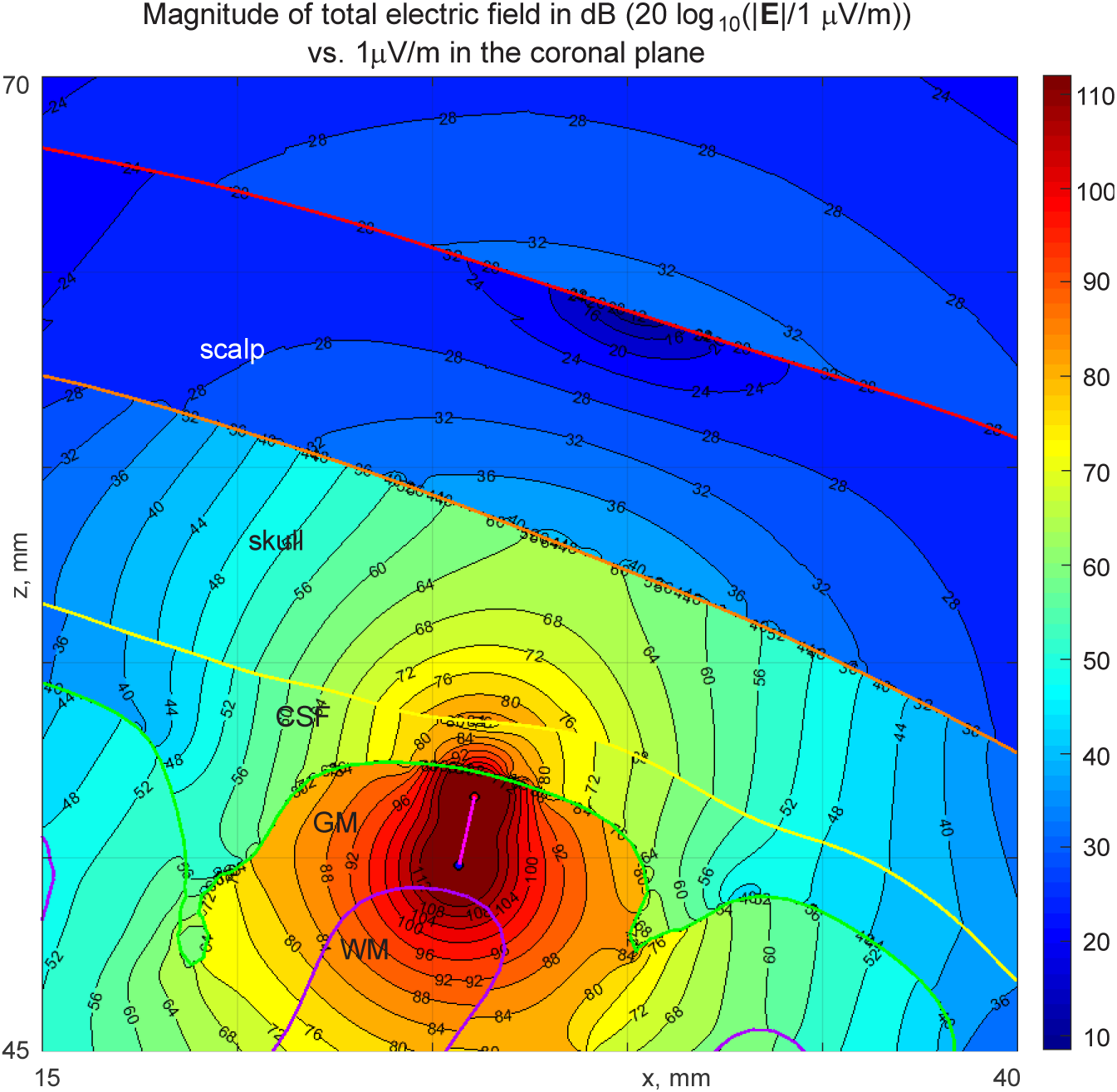
Electric field magnitude distribution in the immediate vicinity of the 1.8 mm long EEG dipole in Fig. 6c. The same logarithmic scale is used but with a better spatial resolution.

### 3.5. A cortical equivalent dipole layer

A spatially extended cortical equivalent dipole layer (He, Yao, and Lian, 2002; Murakami & Okada, 2006; Murakami & Okada, 2015) was simulated using the same high-resolution model as in the previous section. For comparison purposes, this extended source was crated around the single dipole from the previous example. To generate the source, we select all triangular facets of the white matter shell within the distance of 10 mm from the original dipole and assign to every such facet a 1.6 mm long dipole. All the individual dipoles were directed along the outer normal vectors of the white matter facets and are directed outwards as shown in Fig. 8a,b. The distance of each current dipole from the white matter shell was 0.2 mm. Fig. 8b depicts the corresponding dipole distribution to scale. The number of individual dipoles in the present example is ~2,000 although the numbers as large as 20,000 were also tested.

**Fig. 8.**
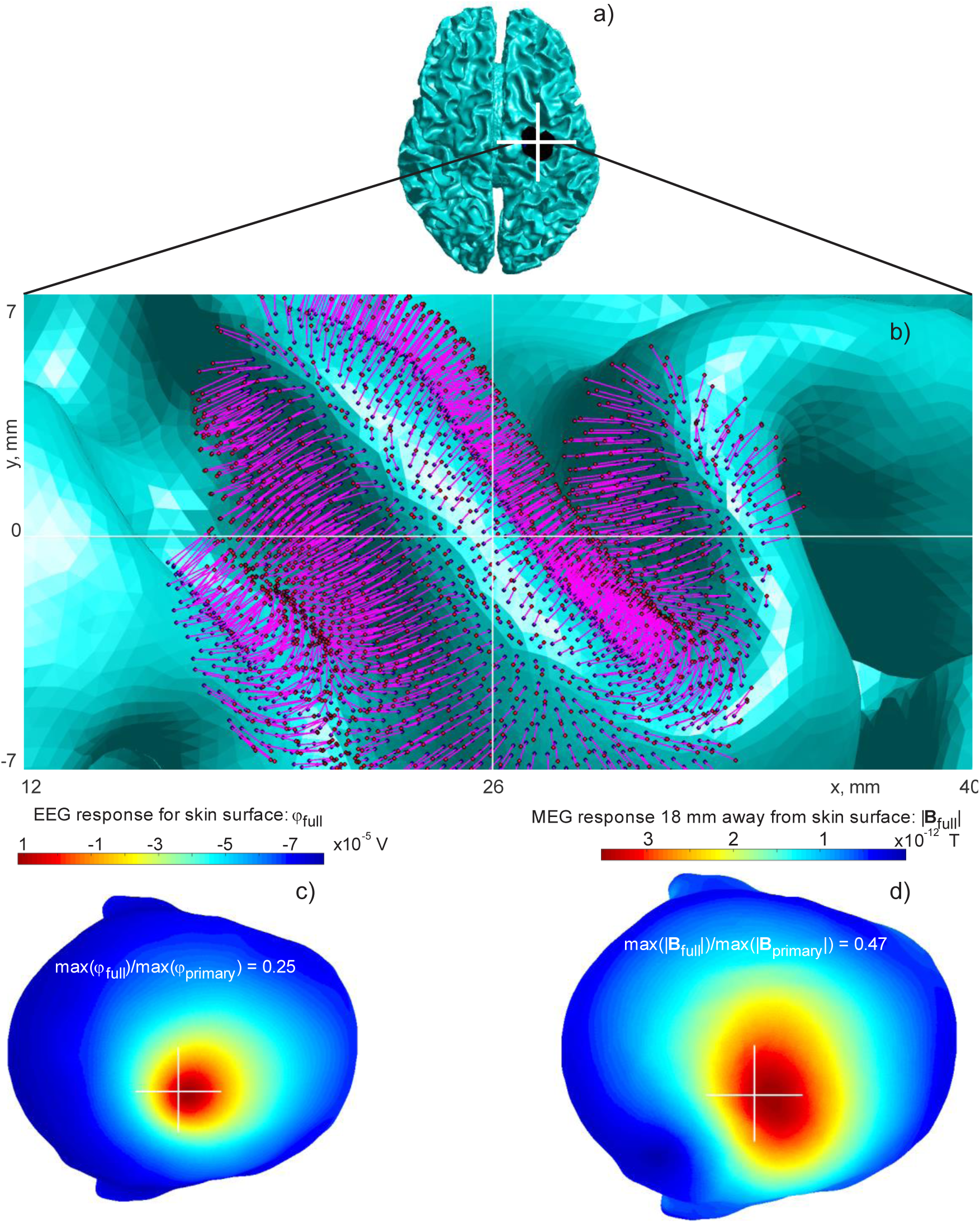
EEG/MEG responses of a small-scale cortical dipole layer computed via BEM-FMM. a), b) Problem geometry;c) The electric potential on the skin surface; d) The magnitude of the total magnetic field 18 mm away from the skin surface. The center of the white cross coincides with projection of the geometrical center of the layer onto a transverse plane.

The Okada-Murakami constant of *q*_0_ = 1 nA ∙ m/mm^2^ (Murakami & Okada, 2006; Murakami & Okada, 2015), also confirmed in a recent MRI study (Sundaram et al., 2016), is used to obtain a realistic current dipole density across the source region. When the dipole length is *d* and the area of the dipole layer is *A*, an expression for the dipole current *I*_0_ follows from *q*_0_ = *I*_0_*d*/*A*, which yields *I*_0_ = *q*_0_*A*/*d*. The total area of the cortical dipole layer is 360 mm^2^. The simulation times are the nearly same as for the single dipole illustrating another computational advantage of the BEM-FMM approach.

The convergence rate is initially very fast but it slows down after the relative residual reaches 10^−3^. Therefore, we restrict ourselves to this value and will discuss this problem later. Fig. 8c shows the resulting surface electric potential on the skin surface and Fig. 8d shows the magnitude of the total magnetic field 18 mm away from the skin surface. A quite significant change in the MEG response pattern is observed as compared to the case of the single dipole located at the cluster center. This is likely due to the presence of multiple horizontal dipoles whose contributions might add up. As to the EEG response pattern, it becomes a bit wider and spatially shifted, but its shape remains nearly the same as for the single dipole.

### 3.6. Conductivity geometries with surface junctions: a head model with a skull opening

Certain problems including modeling of infant MEG/EEG require taking into account more complex anatomical features such as skull openings (fontanels and sutures) that manifest themselves as junctions between the conductivity boundaries. An example with surface junctions in the volume conductor geometry is illustrated in Fig. 9. An adult-sized triangular surface-based head model for subject #101309 from the Population Head Model Repository (Lee et al., 2016; IT’IS Foundation 2017) originated from Connectome Project (Van Essen et al., 2012) was adopted as a starting point The original model was scaled down in order to reflect the average size of a new born head (CDC Growth Charts, 2000).

**Fig. 9.**
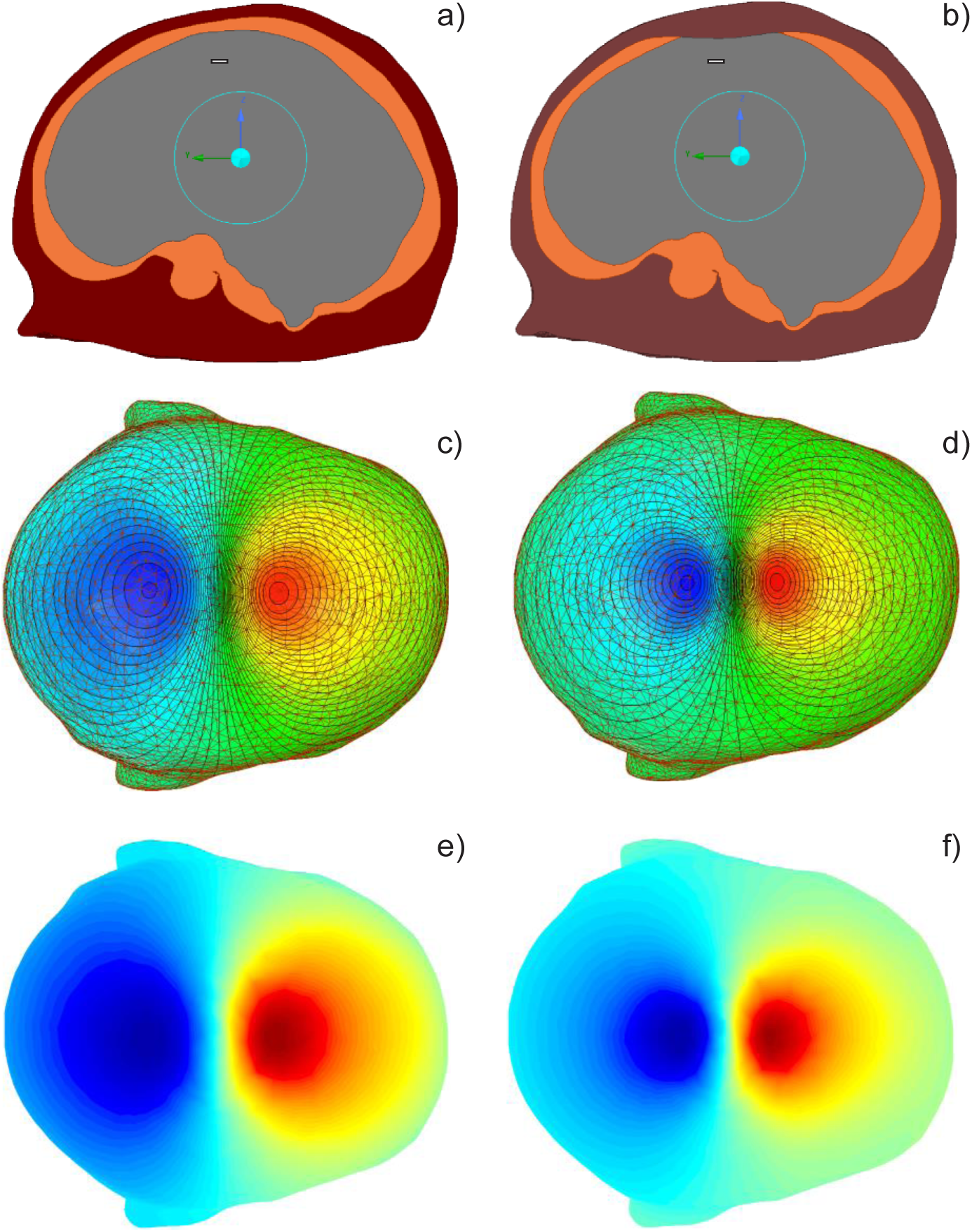
Simplified head geometry without a) and with b) a fontanel; c), d) Voltage distributions on the skin surface obtained with ANSYS Maxwell 3D and normalized to the maximum voltage difference from Table 3; e), f) Voltage distribution on the skin surface obtained via BEM-FMM and normalized to the maximum voltage difference from Table 3.

To mimic the anterior fontanel, the outer skull surface was deformed inwards following a raised cosine elliptical profile with the semi-major axis of 14 mm and the maximum depth of 7 mm. After that, the model was decimated to approximately 10,000 facets per tissue boundary and smoothed using surface-preserving Laplacian smoothing (Vollmer, Mencl, and Müller, 1999a,b). The new boundary surfaces were created by Boolean subtraction of the CSF/brain shell from the deformed skull shell that resulted in a 2-manifold closed skull surface (with the correct model of fontanel) while the intersection of the deformed outer skull shell and the CSF/brain resulted in a new 2-manifold CSF/brain shell. These operations were performed both in ANSYS^®^ Maxwell 3D Electromagnetics Suite 2019 R1 commercial FEM software and within the MATLAB BEM mesh processing environment documented in (Makarov, Noetscher, and Nazarian, 2016).

**Table 3.** Potential/voltage differences on the skin surface in Fig. 9 using ANSYS first-order FEM and BEM-FMM, respectively. A tangential current dipole with the moment *Q* = 1 μA ∙ m is considered.

| Method | Without fontanel | With fontanel |
| --- | --- | --- |
| ANSYS<sup>®</sup> 3D FEM | Max voltage diff.: 0.856 mV | Max voltage diff.: 1.607 mV |
| BEM-FMM, rel. res.=1e-3 | Max voltage diff.: 0.852 mV | Max voltage diff.: 1.542 mV |
| Deviation of maximum difference (FEM vs. BEM-FMM) | 0.5% | 4.0% |
| Average potential error (FEM vs. BEM-FMM) over the entire skin surface following Eq.(16) | 9.6% | 17.9% |

To focus on the effects of the skull opening in the model, only three most essential tissue compartments were kept in the processed model: skin (or scalp) with the conductivity of 0.200 S/m, deformed skull with the conductivity of 0.0132 S/m, and the deformed CSF/brain shell which encloses all brain compartments and which was therefore assigned average newborn brain conductivity of 0.400 S/m. Fig. 9b shows the model thus obtained versus the control model with an intact skull (Fig. 9a). Importantly, all three surfaces (scalp, skull, CSF/brain) are still 2-manifolds and thus closed and orientable. The only global requirement is that the conductivities *inside* the surfaces need to be uniquely defined. We should note that these conditions are satisfied for all practical cases that stem from extracting boundary surfaces of volumetric objects, so the method is completely general. However, for each triangle, the local conductivity contrast (conductivity value on each side of the triangle as defined by its normal vector) needs to be uniquely defined. Therefore, the duplicates of the triangles that are shared by CSF/brain and skull surfaces were identified and removed prior to the execution of the BEM-FMM algorithm.

A tangential electric dipole with the dipole moment *Q* = 1 μA ∙ m (dipole length is 5 mm and source strength is 0.2 mA) was inserted into the brain volume at the distance of 36 mm from origin as shown in Fig. 9a,b. The distance of the dipole from the brain surface was 6.9 mm.

The surface electric voltage was evaluated everywhere on the skin surface using ANSYS^®^ Maxwell 3D FEM software (Electromagnetics Suite 2019 R1, DC Conduction solver module) with 12 adaptive mesh refinement passes and with the final tetrahedral meshes approaching 3 × 10^6^ tetrahedra. Fig. 9c shows the voltage map computed for the head without the skull opening while Fig. 9d is the same results with the skull opening. Both voltage maps are normalized to the maximum voltage differences given in Table 3. The effect of the fontanel is clearly visible on the potential maps. Furthermore, there is a clear agreement between the BEM-FMM computations and ANSYS^®^ FEM results that is illustrated in Fig. 9e,f, respectively. To quantify this, Table 3 lists the maximum voltage differences observed using both methods and the average relative error between the two solutions following Eq. (16). A higher average error for the surface potential may be explained by a non-equivalent modeling of the dipole in ANSYS Maxwell. It is evident from Table 3 that the fontanel has a significant influence on EEG response: the voltage difference in the simulated case nearly doubles.

On the other hand, the MEG responses computed via the BEM-FMM for the same problem and shown in Fig. 10a,b, respectively, do not indicate significant differences. This is in stark contrast to the EEG responses for the horizontal dipole shown in Fig. 9 and for the identical vertical dipole shown for completeness in Fig. 10c,d, respectively. In the last case, we use the log-modulus transformation (John and Draper, 1980)

**Fig. 10.**
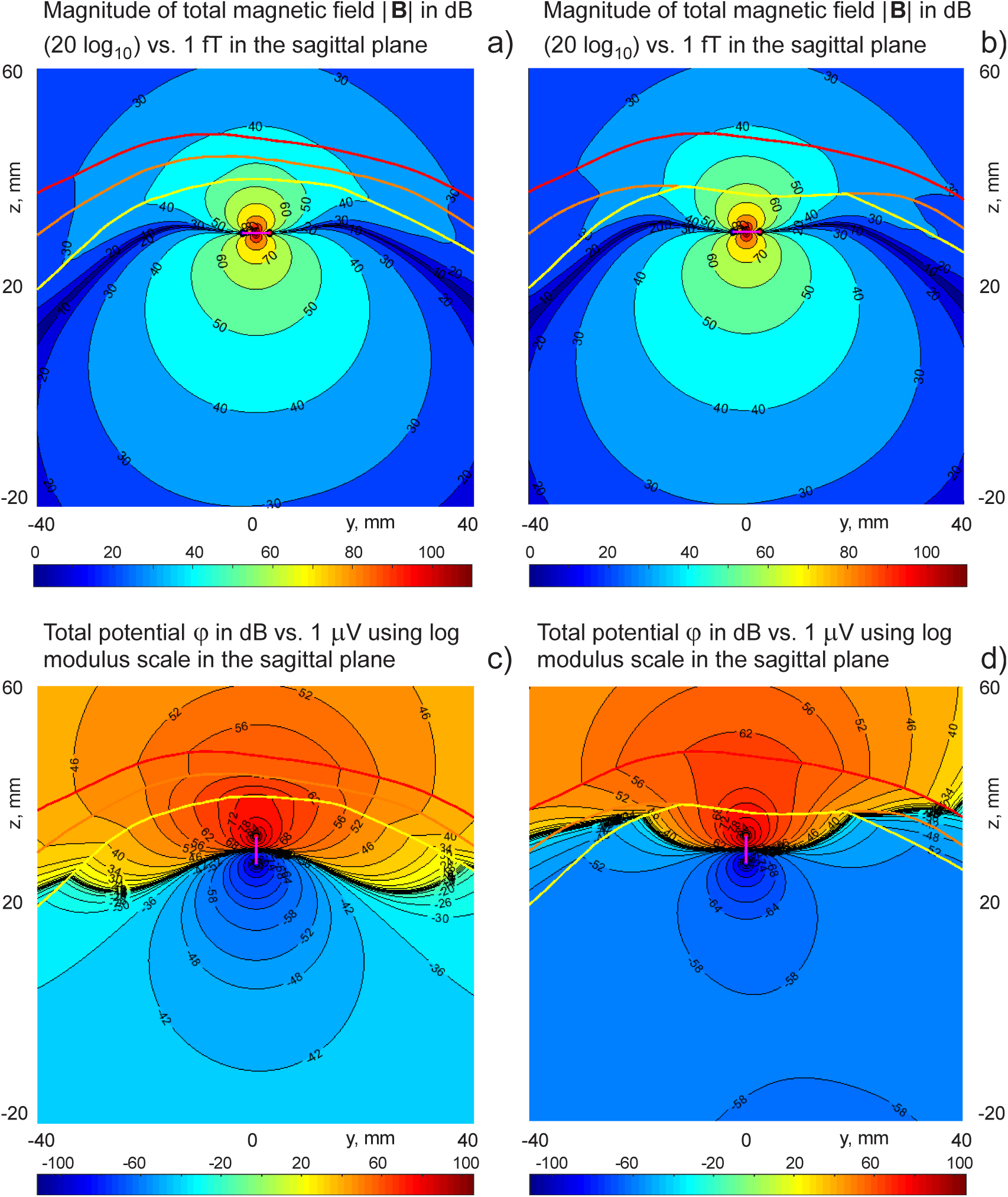
a), b) Distributions of the magnetic field magnitude for the tangential dipole in the sagittal plane obtained via BEM-FMM; c), d) Distributions of the electric potential for the radial dipole in the sagittal plane obtained via BEM-FMM.

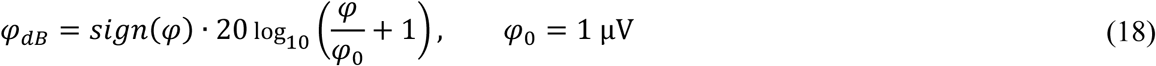

to display large variations of the signed electric potential. In Fig. 10c,d, the maximum on-skin voltage changes from +0.98 mV (no skull opening) to +1.87 mV (with skull opening).

## 4. Discussion and Conclusion

We presented a general computational framework for high-resolution computational modeling of neuroelectromagnetic fields. The method is based on a BEM formulation with several distinct features that allow efficient utilization of the FMM acceleration and is related to our previously developed solver for TMS (Makarov, Noetscher, Raij, and Nummenmaa, 2018; Htet et al., 2019). The presented method is equally well suited for modeling extracranial and intracranial fields and the possible applications include high-resolution modeling of EEG, MEG, ECoG (electrocorticography), and LFP (local field potential) recordings.

In the most demanding clinical evaluations, such as presurgical mapping in epileptic patients, the initial evaluation that starts with EEG and/or MEG is followed by direct recordings with subdural (ECoG) or intraparenchymal depth electrodes. However, despite its vastly improved SNR, ECoG provides only indirect information of the majority of underlying cortical areas, i.e., cortical sulci. In particular, the peaks in ECoG patterns for tangential sources do not coincide with the location of the source. On the other hand, the EEG depth electrodes provide only very limited sparse coverage. Therefore, in principle, the same procedures that are developed for MEG/EEG source modeling could also be applied to greatly improve the value of invasive recordings, which contain more detailed spatial information but the locations of the current sources still need to be computationally inferred from the electric/magnetic field data. Unfortunately, due to the inherent limitations, most crucially due to the surgical penetrations to the skull, no widely accepted application for source modeling for both invasive and non-invasive EEG data still exist.

Resolving the sources of human brain electrophysiological activity poses a significant challenge, specifically in infants or neurological patients whose skull is not intact. The BEM-FMM framework allows handling the most general tissue geometries including junctions between compartments such as skull openings for infants.

The major advantage of the BEM-FMM EEG/MEG modeling approach established in this study is its high speed while maintaining a low computational error and high model resolution as was demonstrated by comparison with the FEM solvers. Furthermore, the method is well suited for a large number of dipoles; the dipoles are also permitted to have a finite length, which makes the method well suited for modeling sources of laminar LFP recordings arising from synaptic activities at different cortical layers. However, the present formulation of the BEM cannot describe anisotropic tissue conductivity, *e.g.*, in the white matter.

We note that a new FMM library is anticipated to be released soon that will enable vectorized computation of multiple dipole distributions for the same head model, resulting in significant reduction in computational cost (L. Greengard, private communication Jan. 10/19). In addition, since the original underlying FMM is using complex arithmetic, the present numerical approach equally well operates with complex dipoles including not only location, magnitude, and direction, but also a phase; it may generate complex electromagnetic fields for each recording channel if desired. Alternatively, reverting to real-valued implementation of the FMM, a factor of two speed-up would be immediately attainable.

Despite significant advantages quantified in this study, the BEM-FMM algorithm is not without its limitations. The FMM part of the BEM-FMM algorithm is not trivial from implementation viewpoint; therefore a general-purpose library is used that may not be ideal for all potential applications. Furthermore, the largest gains in computational efficiency are obtained for problems where high-resolution models are desired. The BEM part of the algorithm relies upon choosing some parameters manually (the number of neighboring triangles for which analytical integration is applied) in order to obtain a good convergence.

An open numerical problem to date is selection of an appropriate stopping criterion of the iterative solution as well as improving its convergence. The relative residual alone is not an entirely adequate measure of the solution convergence to the true result. An initially fast convergence rate of the GMRES method for a realistic head model may saturate, which may indicate that the solution starts to deviate from the true result. We also note that the present method depends on the surface mesh quality and seems to work best for large-scale high-quality smooth manifold meshes. The above-mentioned limitations are shared by many iterative solvers and their practical significance is expected to be small. More research is required to establish standard convergence criteria as well as to quantify the numerical robustness of the results across meshes of varying quality.

We conclude that the presented BEM-FMM approach has high potential to become a standard computational tool for high-resolution forward modeling of electric and magnetic fields of electrophysiological origin.

## Acknowledgements

The authors wish to thank Dr. Leslie Greengard of the Courant Institute of Mathematical Sciences, New York, NY for multiple insightful discussions and support in our utilization of the FMM library. This work has been partially supported by the National Institutes of Health under award numbers R00EB015445, R44NS090894, R01MH111829, R01NS104585, and R01EB022889.

